# Temporal structure of postural sway reveals altered cortical–postural coupling in aging and stroke: insights from nonlinear dynamics and state-space analysis

**DOI:** 10.64898/2026.07.16.738688

**Authors:** Armin Hakkak Moghadam Torbati, Pierre Cabaraux, Thomas Legrand, Scott Mongold, Nicolas Yanguma Munoz, Esranur Yildiran Carlak, Antonella Iannotta, March Vander Ghinst, Gilles Naeije, Lousin Moumdjian, Mathieu Bourguignon

**Affiliations:** Laboratory of Functional Anatomy, Faculty of Human Motor Sciences, Universite libre de Bruxelles (ULB), Brussels, Belgium; Laboratoire de Neuroanatomie et Neuroimagerie translationnelles UNI– ULB Neuroscience Institute, Universite libre de Bruxelles (ULB), Brussels, Belgium; Department of Neurology, Hôpital Universitaire de Bruxelles, CUB Hôpital Erasme, Université libre de Bruxelles (ULB), Brussels, Belgium; School of Electrical and Electronic Engineering, University College Dublin (UCD), Dublin, Ireland; Insight Research Ireland Centre for Data Analytics, University College Dublin, Dublin, Ireland; Service d’ORL et de chirurgie cervico-faciale, HUB Hôpital Erasme, Université libre de Bruxelles (ULB), Brussels, Belgium; Centre de Référence Neuromusculaire, Department of Neurology, CUB Hôpital Erasme, Université libre de Bruxelles (ULB), 1070Brussels, Belgium; REVAL Rehabilitation Research Center, Faculty of Rehabilitation Sciences, Hasselt University, Hasselt, Belgium; WEL Research Institute Avenue Pasteur 6, 1300 Wavre, Belgique

**Keywords:** Corticokinematic Coherence, EEG, CoP, Nonlinear Features, Balance Control, Postural Sway, State-space

## Abstract

**Background:** Balance maintenance in humans is not only a mechanical process, but it also relies on continuous interactions between cortical activity and body dynamics. Alterations in postural sway are commonly observed in aging and stroke and are frequently used to assess balance impairment. However, similar balance deficits do not necessarily reflect similar underlying sensorimotor control mechanisms. Therefore, investigating brain–body interactions and their relationship to characteristics of postural behavior may provide deeper insights into the neural processes underlying balance dysfunction in these populations.

**Objective:** To determine whether brain–body coupling is associated with characteristics of postural behavior captured by the temporal organization of postural fluctuations beyond conventional magnitude-based measures of postural sway, and whether these relationships differ between stroke survivors, healthy older adults, and young adults.

**Methods:** EEG and center-of-pressure (CoP) signals were recorded simultaneously in stroke survivors (n = 12), healthy older adults (n = 18), and young controls (n = 17) during quiet standing under 4 different manipulated sensory conditions. Sway-based corticokinematic coherence (CKC) as well as linear and nonlinear features (sample entropy, SE; fractal dimension, FD) of CoP were extracted. Linear mixed-effects model assessed associations between features and CKC, and model performance was compared using Akaike Information Criterion. Multidimensional state vectors were constructed from CKC, linear and nonlinear CoP features, and Euclidean distances between consecutive states in the standardized feature space were computed to quantify condition-dependent transitions in brain–body control organization.

**Results:** Nonlinear features showed significant, group– and feature-dependent associations with CKC in the mediolateral direction, driven by significant SE and FD effects in the stroke group and an SE effect in the older group, while no significant associations were observed in the young group. Including nonlinear features in baseline models containing only linear CoP features significantly improved model fit. CKC alone showed low classification performance (AUC 50 to 65), whereas combining CKC with linear and nonlinear features improved group discrimination (AUC up to 0.86). State-space transition analysis revealed larger condition-dependent transitions in stroke participants compared with healthy older adults, particularly going from eyes open to eyes closed when standing on foam.

**Conclusion:** Brain–body coupling during standing may be understood more comprehensively by factoring in the temporal structure of fluctuations rather than their amplitude alone. These findings support the use of nonlinear dynamical features, combined with CKC, as potential markers of balance impairment.

## 1. Introduction

Balance maintenance in humans is not only a mechanical process, but it also relies on continuous interactions between cortical activity and body dynamics [1]. These interactions are reflected as unavoidable postural sway during quiet standing [2]. Although postural sway naturally varies across individuals, its alterations have been widely used to characterize balance impairments associated with aging and neurological disorders because they provide valuable information about the underlying sensorimotor control mechanisms [3]. For example, stroke often disrupts sensorimotor function, which leads to increased sway and subsequently elevated fall risk [4]. Similarly, elderly individuals often experience a gradual decline in sensorimotor function, which negatively affects postural stability [5]. Although stroke and aging can produce comparable balance impairments, similar behavioral outcomes do not necessarily imply similar underlying control processes. Therefore, understanding how brain–body interactions affect balance control in these populations could provide more insights into the mechanisms underlying balance impairment and postural dysfunction.

The interaction between brain activity and postural sway is commonly investigated through the simultaneous recording of neuroimaging signals, such as electroencephalography (EEG), along with center-of-pressure (CoP) measurements. In this context, sway-based corticokinematic coherence (CKC) is a promising approach used to quantify the interaction between cortical activity measured with EEG and CoP fluctuations using coherence analysis [6]. Previous studies have shown that CKC is strongest over sensorimotor cortical regions and modulated by task demands. Moreover, CKC appears to reflect the functional coupling within the sensorimotor loop, encompassing both efferent motor commands and afferent sensory feedback [7–9]. However, CKC primarily quantifies the strength of synchronization between cortical activity and postural sway. Although this measure provides information about the degree of cortico-postural interaction, it does not reveal which characteristics of postural behavior are associated with that interaction. In particular, CKC alone cannot determine whether stronger brain–body coupling is associated only with the magnitude of postural sway or also with the temporal organization of postural fluctuations.

To facilitate the behavioral interpretation of CKC, previous studies have mainly related CKC to conventional magnitude-based measures of postural sway, such as CoP standard deviation and mean CoP velocity, demonstrating significant associations between cortico-postural coupling and these linear descriptors of balance performance [8–10]. Nevertheless, to our knowledge, no study has directly related CKC to nonlinear features of postural sway to examine how cortico-postural interaction relates to the temporal structure and complexity of postural sways. This gap is critical because the temporal organization of sway may reflect underlying brain-body control strategies rather than its magnitude [11]. Two signals with comparable amplitude can differ substantially in their temporal structure, indicating distinct control mechanisms that are not captured by conventional amplitude– or variability-based metrics [11, 12]. In this context, nonlinear measures such as sample entropy (SE) and fractal dimension (FD) provide a principled method to quantify the irregularity and complexity of postural sway across time [13].

Accordingly, this study aimed to determine whether cortico-postural coupling is associated with the temporal organization of postural sway beyond its previously reported associations with conventional magnitude-based measures and whether these associations differ among stroke, healthy older adults, and young controls. To this end, we examined the relationship between CKC and nonlinear features of CoP signals across stroke survivors, healthy older adults, and young controls under varying sensory conditions. Balance control depends on the integration and reweighting of visual, vestibular, and somatosensory information; therefore, varying sensory conditions may reveal how cortical–postural interactions adapt to changing sensory demands [14]. In addition, while feature-specific analyses can identify individual associations between CKC and postural sway characteristics, they cannot reveal how neural and behavioral variables jointly reorganize as an integrated brain–body control system. Therefore, we additionally explored the evolution of multidimensional brain–body states across sensory challenges. We hypothesized that associations between CKC and nonlinear sway features would be stronger in stroke survivors and healthy older adults than in young adults because impaired automatic balance control may increase the dependence of cortical– postural coupling on the temporal organization of sway. If so, it could provide informative markers of balance impairment and offer insights into how sensorimotor control is differentially altered in aging and stroke. Such findings may provide a foundation for developing more sensitive and mechanistically informed biomarkers of balance dysfunction grounded in brain–body interaction dynamics.

## 2. Methods and materials

This study reanalysed previously published data [8, 9]. An overview of the experimental design, signal processing, and analytical pipeline is presented in Fig. 1.

**Fig. 1.**
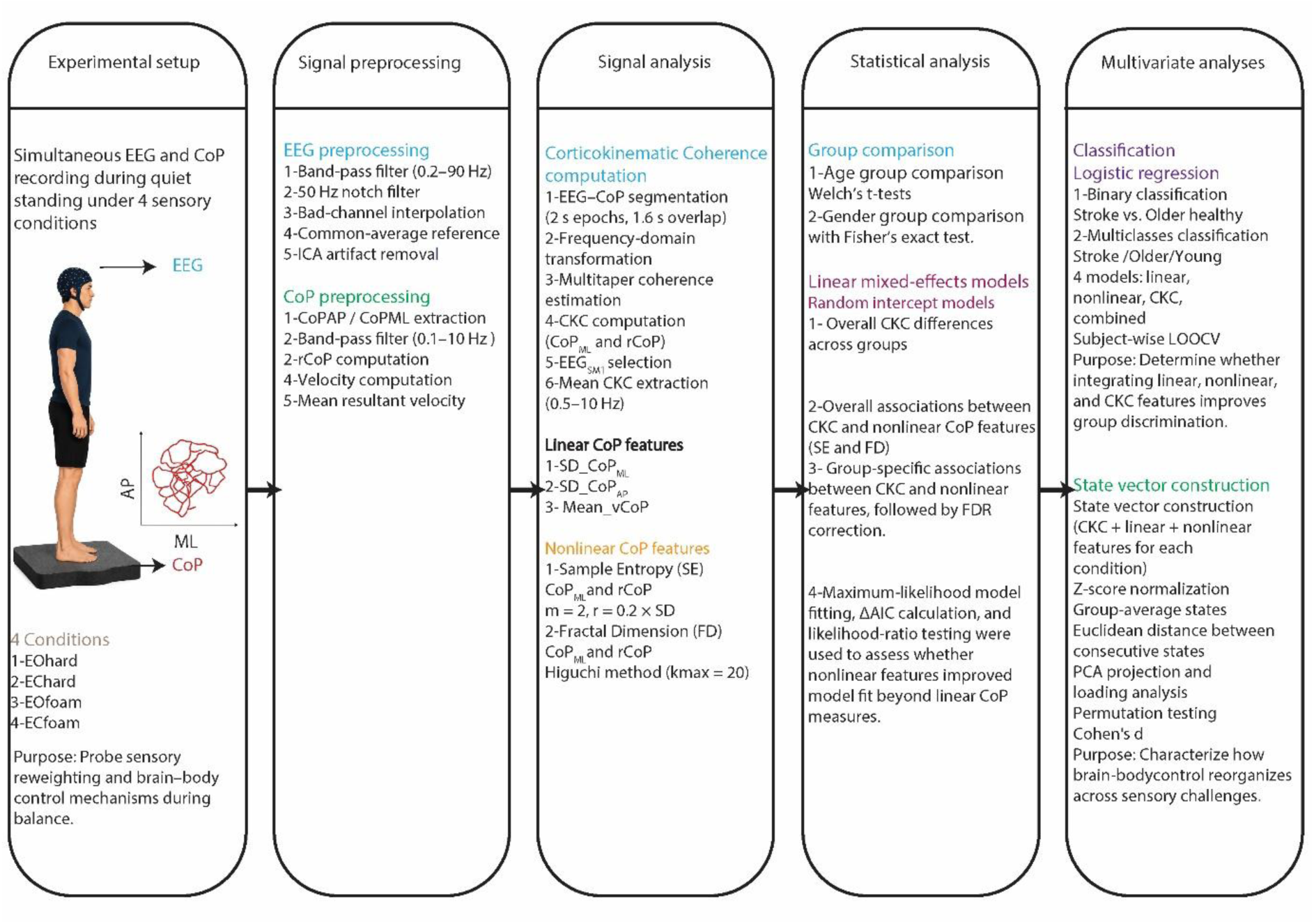
Overview of the experimental and analytical workflow.

### 2.1. Participants

The study included 12 stroke patients (mean ± SD; age: 67.0 ± 16.6 years; 3 women), 18 healthy older adults (70.8 ± 6.9 years; 10 women), and 17 healthy young subjects (25.5 ± 2.2 years; 6 women). Stroke participants exhibited relatively mild functional impairment, as reflected by high Berg Balance Scale scores (mean ± SD; 51.5 ± 7.0) and Functional Ambulation Category scores (mean ± SD; 4.7 ± 0.6). Lesions were heterogeneous in both location and etiology, including ischemic and hemorrhagic strokes affecting cortical, subcortical, cerebellar, and brainstem regions. Overall, the clinical characteristics indicate that they were independently ambulatory and capable of performing the standing balance tasks required in the study. Healthy volunteers were excluded if they reported a history of neurological, muscular, or vestibular disorders. The study protocol was approved by the Erasme University Hospital Ethics Committee (CCB: B4062021000323, Brussels, Belgium), and all procedures adhered to the relevant institutional guidelines. All participants provided written informed consent prior to enrollment.

### 2.2. Experimental protocol

Participants across all three groups performed a series of balance tasks under four different sensory conditions designed to systematically manipulate sensory input. These conditions included bipedal stance (1) on a hard (firm) surface with eyes open (EOhard), (2) on a hard surface with eyes closed (EChard), (3) on a foam surface with eyes open (EOFoam), and (4) on a foam surface with eyes closed (ECfoam). Each condition lasted for 5 minutes and the resulting 4 recordings blocks were performed in a randomized order. To maintain consistency, all participants stood barefoot with their feet parallel at shoulder-width, arms at their sides, without speaking, and with eyes fixed on a wall-mounted cross. An assessor stood directly behind them to ensure safety and to provide assistance in the event that a fall occurred; however, no physical assistance was needed during the trials. To minimize the impact of fatigue, participants were closely monitored for self-reported fatigue and provided with mandatory rest periods between trials.

### 2.3. Recordings

EEG was recorded during the postural balance tasks with 64 electrodes (ANT Neuro, Hengelo, Netherlands). Electrodes, embedded in a cap, were arranged according to the 10/20 system. Electrolyte gel was applied to ensure electrode impedance was below 20 kΩ, and the signal was referenced to CPz. All EEG data were sampled at 1000 Hz. Simultaneously, a force plate (AccuSway-O, AMTI, Watertown, MA, USA) recorded ground reaction forces and moments at 1000 Hz.

### 2.4. Preprocessing

EEG and force plate recordings were analyzed with MATLAB (MathWorks, Natick, MA, USA, R2024b). Investigation of the raw data across all participants showed that mastoid electrodes (M1 and M2) exhibited high-amplitude artifacts due to poor or unstable skin–electrode contact. Consequently, these electrodes were excluded, and only the remaining channels were retained for subsequent analysis.

EEG signals were band-pass filtered between 0.2 and 90 Hz and additionally processed with a 50 Hz notch filter using a fourth-order Butterworth filter. Bad channels were detected following the criteria proposed by Bidgely-Shamlo et al. [15]. Signals from these channels were then reconstructed via interpolation using neighboring electrode signals [16]. Subsequently, the EEG data were re-referenced to the common average. Next, 20 independent components were extracted using Fast Independent Component Analysis (ICA) [17] to further isolate physiological artifacts. Components associated with cardiac activity, eye blinks, and eye movements were visually identified, and their contributions were removed from the full-rank data which was reconstructed by the mixing matrix. Finally, data points where EEG amplitude exceeded the mean by more than 5 standard deviations were marked as bad.

### 2.5. Data analysis

#### 2.5.1. Center of Pressure (CoP) computation

The CoP time-series, including the antero-posterior (CoP_AP_) and mediolateral (CoP_ML_) components, were band-pass filtered between 0.1 and 10 Hz using a fourth-order Butterworth filter. The excursion (rCoP) was calculated as the Euclidean norm of CoP_AP_ and CoP_ML_ at each time point. CoP velocities in the antero-posterior and mediolateral directions were obtained as the first derivative of the CoP time-series. The resultant CoP velocity was then computed as the Euclidean norm of these velocity components at each time point, and its mean value was used as the mean resultant CoP velocity (Mean_vCoP).

#### 2.5.2. Sway-based CKC computation

Sway-based CKC was evaluated through coherence analysis, which is an extention of the Pearson correlation coefficient to the frequency domain. This method quantifies the coupling between two signals, providing a value between 0 (no linear relationship) and 1 (perfect linear relationship) at each frequency [18]. In practice, EEG and CoP signals were segmented into overlapping 2-second epochs with a 1.6-second overlap, yielding a frequency resolution of 0.5 Hz. Analysis was restricted to four electrodes that typically overlie the lower limb sensorimotor area (SM1) (Cz, C3, C4, Fz), and based only on epochs more than 1 s away from bad samples. To maintain consistency across experimental conditions, epochs were removed so that each participant had a comparable number of artifact-free epochs in all conditions (534 ± 6, mean ± SD). The excluded epochs corresponded to the final ones of the last recording block.

The remaining epochs were subsequently transformed into the frequency domain using the fast Fourier transform and combined to compute coherence spectra between each of the two selected CoP features (CoP_ML_, and rCoP) and each of the retained EEG channels. This was done according to the approach described by Halliday et al. [18] and using a multitaper method (with 3 orthogonal Slepian tapers, resulting in a spectral smoothing of 1.5 Hz) to estimate power– and cross-spectral densities [19].

We chose rCoP because previous sway-based CKC studies consistently identified robust between cortical activity and rCoP, and because rCoP appears to reflect the corrective output component of the postural control loop. Specifically, previous work showed that changes in cortical activity tend to precede changes in rCoP, suggesting that rCoP is closely related to the motor commands used to regulate posture. In contrast, sway velocity appears to be more strongly associated with sensory-feedback processing, where postural fluctuations are first monitored and then represented in cortical activity. Therefore, rCoP may provide a more direct window into cortical contributions to postural regulation rather than sensory monitoring alone [8, 10]. In addition, CoP_ML_ was included since stroke patients often exhibit postural asymmetry [20], potentially revealing differences in cortical involvement associated with asymmetrical weight distribution during balance control. For subsequent analyses, only the electrode showing the highest CKC, averaged across balance conditions and within the 0.5–10 Hz range, was retained for each feature. This electrode is hereafter referred to as EEG_SM1_. CKC strength was then quantified as the mean coherence value within the 0.5–10 Hz band at the EEG_SM1_ electrode.

#### 2.5.3. Linear feature extraction from CoP

We extracted linear features including the standard deviation of the CoP along the mediolateral (SD_CoP_ML_) and antero-posterior axes (SD_CoP_AP_) and the mean of computed resultant CoP velocity (Mean_vCoP).

### 2.6. Non-linear dynamic feature extraction from CoP

We also extracted 2 nonlinear dynamic features, SE and FD, from CoP_ML_ and rCoP. In all cases, the computation was performed on the preprocessed CoP signals that were fruther resampled to 100 Hz and linearly detrended.

#### 2.6.1. Sample Entropy

SE quantifies the irregularity and unpredictability of a time-series signal. Specifically, it estimates the probability that patterns (vectors) of length m, which are similar within a defined tolerance r, remain similar when the sequence length increases to m+1. Compared to Approximate Entropy, SE reduces bias by excluding self-matches and exhibits greater consistency across different data lengths, although it still requires sufficient data for reliable estimation [21]. Owing to these properties, SE has been widely used in CoP analysis to assess the complexity of postural control.

SE was calculated using an embedding dimension of m = 2, time delay τ = 1 sample, and tolerance r = 0.2 times the standard deviation of the processed signal [22]. Similarity between vectors was assessed using the Chebyshev distance, self-matches were excluded, and SE was computed as −log(A/B), where B and A denote the numbers of matching vector pairs of length m and m + 1, respectively [23]. The length of the time series was sufficient to ensure reliable estimation of SE [24].

#### 2.6.2. Fractal Dimension

FD provides a quantitative measure of the geometric complexity and scaling properties of CoP trajectories. Unlike traditional measures that quantify the magnitude or velocity of sway, FD characterizes how the CoP trajectory occupies space across multiple scales, thereby reflecting the intrinsic organization of variability in postural fluctuations. In quiet standing, CoP signals exhibit fractal-like behavior [25], indicating that postural sway arises from a complex, nonlinear control system rather than from random or deterministic processes. Lower values of FD indicate more regular and constrained trajectories, and higher values reflect increased irregularity and spatial complexity [26].

The Higuchi algorithm was then applied using scales k = 1 to 20. At the analyzed sampling rate of 100 Hz, this corresponds to a maximum temporal interval of 0.20 s. The choice of kmax = 20 was based on previous studies and selected to ensure a balance between temporal resolution and stability of estimation [27]. For each scale, multiple sub-series were constructed, their normalized curve lengths were averaged, and FD was obtained as the slope of the log–log relationship between average curve length and inverse scale. The same kmax value was used for all participants and conditions to ensure consistency of estimation.

### 2.7. Statistical analysis

Age differences between stroke and older healthy groups were assessed using independent two-sample Welch’s t-tests to account for unequal sample size. Differences in sex distribution between these groups were evaluated using Fisher’s exact test.

Linear mixed-effects models were used throughout the study to account for the repeated-measures structure of the data, with observations across the four experimental conditions (EOhard, EOfoam, EChard, and ECfoam) nested within subjects. All analyses were performed separately for CoP_ML_ and rCoP derived features. In all models, subject was included as a random intercept to account for repeated measurements within individuals, and statistical significance of fixed effects was evaluated using ANOVA with significance set at α = 0.05.

To examine whether CKC differed between groups across experimental conditions, a model including group, condition, and their interaction as fixed effects was fitted.

To investigate the relationship between nonlinear CoP features and CKC, initial analyses were performed using the entire dataset. These models included SE, FD, group, and condition as fixed-effect predictors. To determine whether the associations between nonlinear features and CKC differed across groups, an additional model was fitted including group-by-feature interaction terms together with the corresponding SE, FD, group, and condition.

To further investigate whether the observed associations were group-specific, follow-up analyses were conducted separately within each group (stroke survivors, healthy older adults, and healthy young adults). In these models, SE, FD, and condition were included as fixed-effect predictors to assess their associations with CKC. P-values from these follow-up analyses were adjusted using the false discovery rate (FDR) procedure, with significance assessed at an FDR-corrected α = 0.05.

Since the primary objective of the present study was to investigate whether nonlinear features of postural sway are associated with CKC and whether they explain variance beyond conventional linear measures, post hoc analyses and multiple-comparison corrections were applied only to the nonlinear feature analyses.

To determine whether nonlinear features explained variance in CKC beyond that captured by conventional linear measures of postural sway, a model-comparison approach was employed. For each analysis, a baseline model was first constructed containing the classical linear sway descriptors: standard deviation of CoP displacement in the mediolateral (SD_CoP_ML_) and anteroposterior (SD_CoP_AP_) directions, mean CoP velocity (mean_vCoP), and experimental condition as fixed effects. Subject was included as a random intercept. Expanded models were then constructed by adding a single nonlinear feature of interest (SE or FD) to the baseline model while retaining all baseline predictors. Separate comparisons were performed for each nonlinear feature and for each group.

Models were fitted using maximum likelihood (ML), and model fit was first evaluated using the Akaike Information Criterion (AIC), where lower values represent a better balance between goodness-of-fit and model complexity [28, 29]. To quantify the relative improvement in model fit, differences in AIC (Δ*AIC*) were calculated between the baseline linear model and each extended model as:

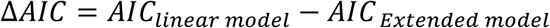

Positive ΔAIC values indicate improved model fit when the nonlinear feature is included [28]. In addition, nested models were formally compared using likelihood ratio tests (LRTs), yielding X^2^ statistics and corresponding p-values. These analyses were used to determine whether inclusion of a nonlinear feature significantly improved model fit beyond that achieved by conventional linear sway measures alone.

### 2.8. Classification Analysis

Classification analyses were performed to discriminate between stroke and older healthy subjects and between older and younger subjects (binary classification), as well as among all three groups (stroke, older healthy, and younger healthy; multiclass classification). In each scenario, four feature sets were used to construct logistic regression models: (i) linear CoP features (SD_CoP_ML_, SD_CoP_AP_, and mean_vCoP), (ii) nonlinear CoP features (SE_CoP_ML_ and FD_CoP_ML_), (iii) CKC, and (iv) a combination of these 3 sets.

To account for repeated measurements, model performance was evaluated using subject-wise leave-one-out cross-validation, in which all observations (conditions) from a given subject were excluded from the training set and used exclusively for testing. Predictions from all observations of a subject were aggregated by averaging the predicted probabilities to obtain a single subject-level score. Features were standardized (z-scored) within each training fold, and the same transformation was applied to the corresponding test data to avoid data leakage.

Logistic regression models were fitted to the training data, and predicted probabilities were obtained for the test observations. For binary classification, performance was evaluated using accuracy, area under the receiver operating characteristic (ROC) curve (AUC), sensitivity, and specificity. For multiclass classification, performance was assessed using overall accuracy and confusion matrices, from which class-specific recall (row-normalized) and precision (column-normalized) were derived for each group.

### 2.9. Brain–body state-space trajectory analysis

Previous analyses examined individual relationships between CKC and specific sway features. However, balance control emerges from the interaction of multiple neural and behavioral processes operating simultaneously. Therefore, examining features individually may overlook how these variables jointly reorganize as a coordinated brain–body control system across sensory challenges. To address this gap, we constructed multidimensional brain–body state vectors combining CKC, linear sway measures, and nonlinear sway measures, and examined how these states evolved across progressively challenging balance conditions.

For each participant and each sensory condition, a multivariate brain–body state vector was constructed. Each state vector included CKC, linear CoP features (SD_CoP_ML_, SD_CoP_AP_, and Mean_vCoP), and nonlinear CoP features (SE and FD). Thus, each participant was represented by four state vectors corresponding to the four sensory conditions.

Before computing distances, all state-space features were standardized using z-score normalization across all participants and conditions. This normalization was applied to ensure that features with different numerical scales contributed comparably to the multivariate distance calculations.

For each participant, the four sensory-condition state vectors were ordered according to the experimental progression of sensory challenge: EOhard → EChard → EOfoam → ECfoam

Euclidean distances were then computed between consecutive state vectors within the standardized feature space. Specifically, three transition distances were calculated: D1, corresponding to the transition from EOhard to EChard; D2, corresponding to the transition from EChard to EOfoam; and D3, corresponding to the transition from EOfoam to ECfoam. These distances quantified the magnitude of brain–body state reconfiguration required between successive sensory conditions.

To visualize the organization of brain–body state trajectories, principal component analysis (PCA) was applied to the standardized state vectors from all participants and conditions. The first two principal components were used only for visualization of group-level trajectories. For each group, mean PC scores were computed for each sensory condition and plotted in the predefined condition order to illustrate the average trajectory of brain–body state evolution across sensory challenge. PCA weights were also extracted to quantify the contribution of each feature to the first two principal components.

To statistically assess group differences in state-space reconfiguration, permutation testing was performed on the subject-level transition distances (D1, D2, and D3). For each pairwise group comparison and each transition metric, the observed difference in mean transition distance between groups was calculated. Group labels were then randomly permuted 10,000 times, and the mean difference was recomputed for each permutation to generate a null distribution. Two-sided permutation p-values were calculated as the proportion of permuted absolute mean differences greater than or equal to the observed absolute mean difference. Cohen’s d was also computed to quantify effect size for each comparison.

## 3. Results

No significant differences were observed between the stroke and older healthy groups in either age (p = 0.46) or sex distribution (p = 0.14).

### 3.1. Corticokinematic Coherence

Fig. 2 presents the distribution of CKC values across groups and experimental conditions for CoP_ML_ and rCoP.

**Fig. 2.**
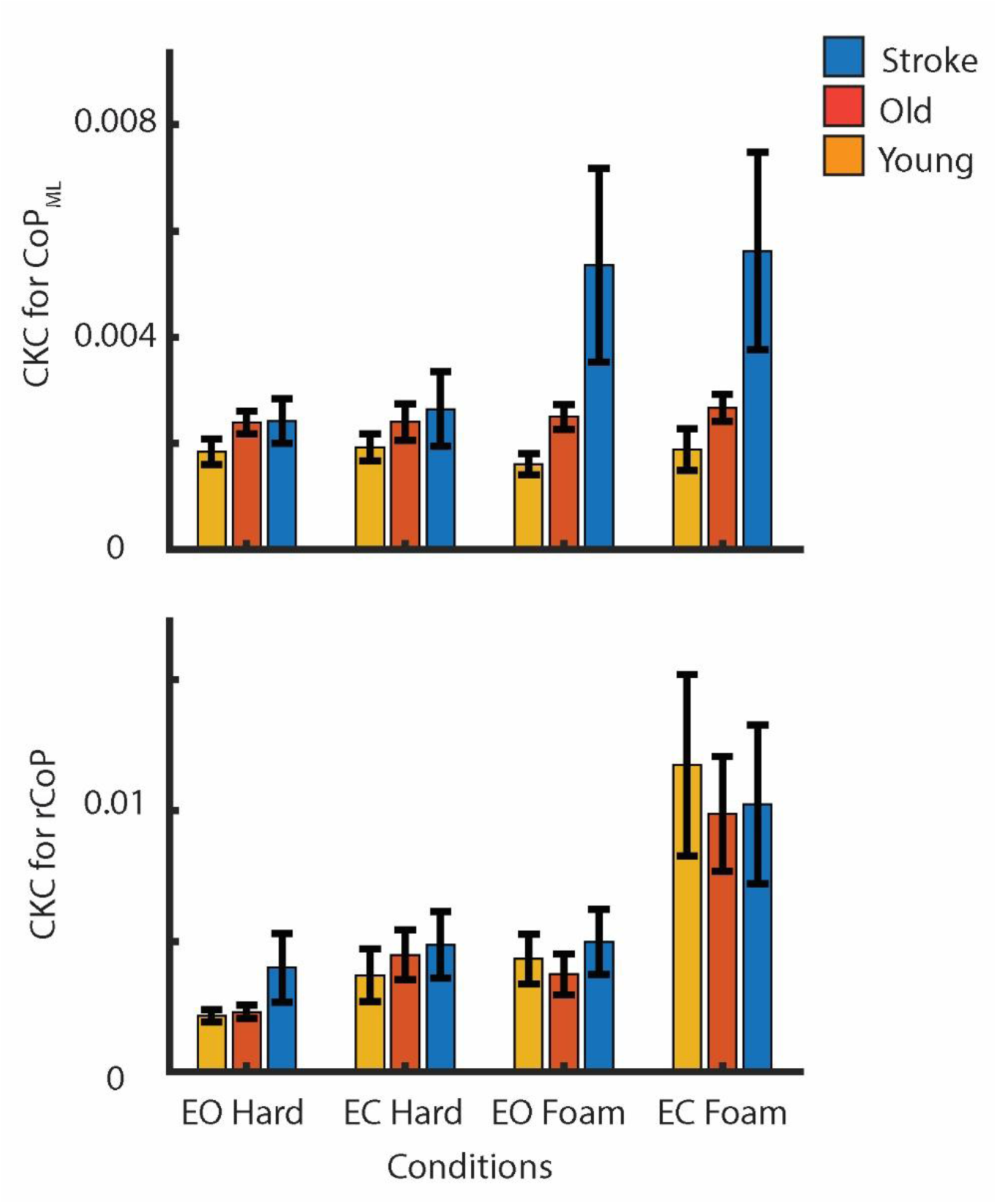
Mean ± SEM of CKC across groups and conditions. (**Upper panel**) CKC for CoP_ML_. (Lower panel) CKC for rCoP.

For CKC for CoP_ML_, no significant main effect of condition was observed (F(3,176)=0.06, p=0.97). A significant main effect of group was found (F(2,176)=8.49, p<0.001), along with a significant group by condition interaction (F(6,176)=2.78, p=0.01).

For CKC for rCoP, a significant main effect of condition was observed (F(3,176)=6.36, p<0.001). In contrast, neither the main effect of group (F(2,176)=0.41, p=0.66) nor the group by condition interaction (F(6,176)=0.32, p=0.92) reached statistical significance.

### 3.2. Nonlinear CoP features and group-specific associations with CKC

The overall relationships between CKC and nonlinear features of postural sway were first examined across all participants before conducting group-specific analyses.

Significant associations were observed between CKC for CoP_ML_ and nonlinear features of postural sway. Specifically, significant effects of both SE (F(1,176)=9.63, p=0.002) and FD (F(1,176)=12.67, p<0.001) were observed. In addition, significant SE × group (F(2,176)=18.86, p<0.001) and FD × group (F(2,176)=20.24, p<0.001) interactions were also observed, indicating that the relationships between CKC and nonlinear features differed across stroke survivors, healthy older adults, and healthy younger adults.

No significant associations were observed between CKC for rCoP and the nonlinear features of postural sway. Specifically, neither SE (F(1,176)=0.03, p=0.86) nor FD (F(1,176)=0.98, p=0.32) showed a significant relationship with CKC.

Group-specific analyses revealed distinct relationships between CKC for CoP_ML_ and nonlinear features of postural sway. In stroke survivors, both SE (F(1,42)=13.08, P_FDR_=0.002) and FD (F(1,42)=15.91, P_FDR_=0.002) were significantly associated with CKC. In healthy older adults, a significant association was observed between SE and CKC (F(1,65)=10.44, P_FDR_=0.004), whereas FD did not reach statistical significance (F(1,65)=1.85, P_FDR_=0.26). In contrast, neither SE (F(1,59)= 0.69, P_FDR_=0.48) nor FD (F(1,59)= 0.01, P_FDR_=0.88) was significantly associated with CKC in healthy younger adults.

### 3.3. Improvement of model performance by nonlinear features

In the CoP_ML_ direction, to determine whether nonlinear features explained variance in CKC beyond conventional linear CoP measures, model fit was compared between baseline models containing linear features only and extended models including either SE or FD. In stroke survivors, the inclusion of both SE (ΔAIC = 4.63, X^2^(1) = 6.63, p = 0.010) and FD (ΔAIC = 7.31, X^2^(1) = 9.31, p = 0.002) improved model fit relative to the baseline model. In healthy older adults, inclusion of SE improved model fit (ΔAIC = 2.07, X^2^(1) = 4.07, p = 0.044), whereas inclusion of FD did not result in a significant improvement (ΔAIC = 1.12, X^2^(1) = 3.12, p = 0.077).

### 3.4. Classification of groups

Table 1 presents the result of the classification analysis of the four models applied to classify stroke versus older healthy subjects. Linear-only and CKC-only models showed moderate performance, with accuracies around 60%; however, their AUC values and, in particular, sensitivities were low (sensitivity = 0.17). In contrast, the nonlinear model demonstrated better evaluation metrics, achieving 77% accuracy, an AUC of 0.73, and improved sensitivity (0.58). The combined model further increased the AUC to 0.80 while maintaining a similar accuracy (Fig. 3, Table 1).

**Fig. 3.**
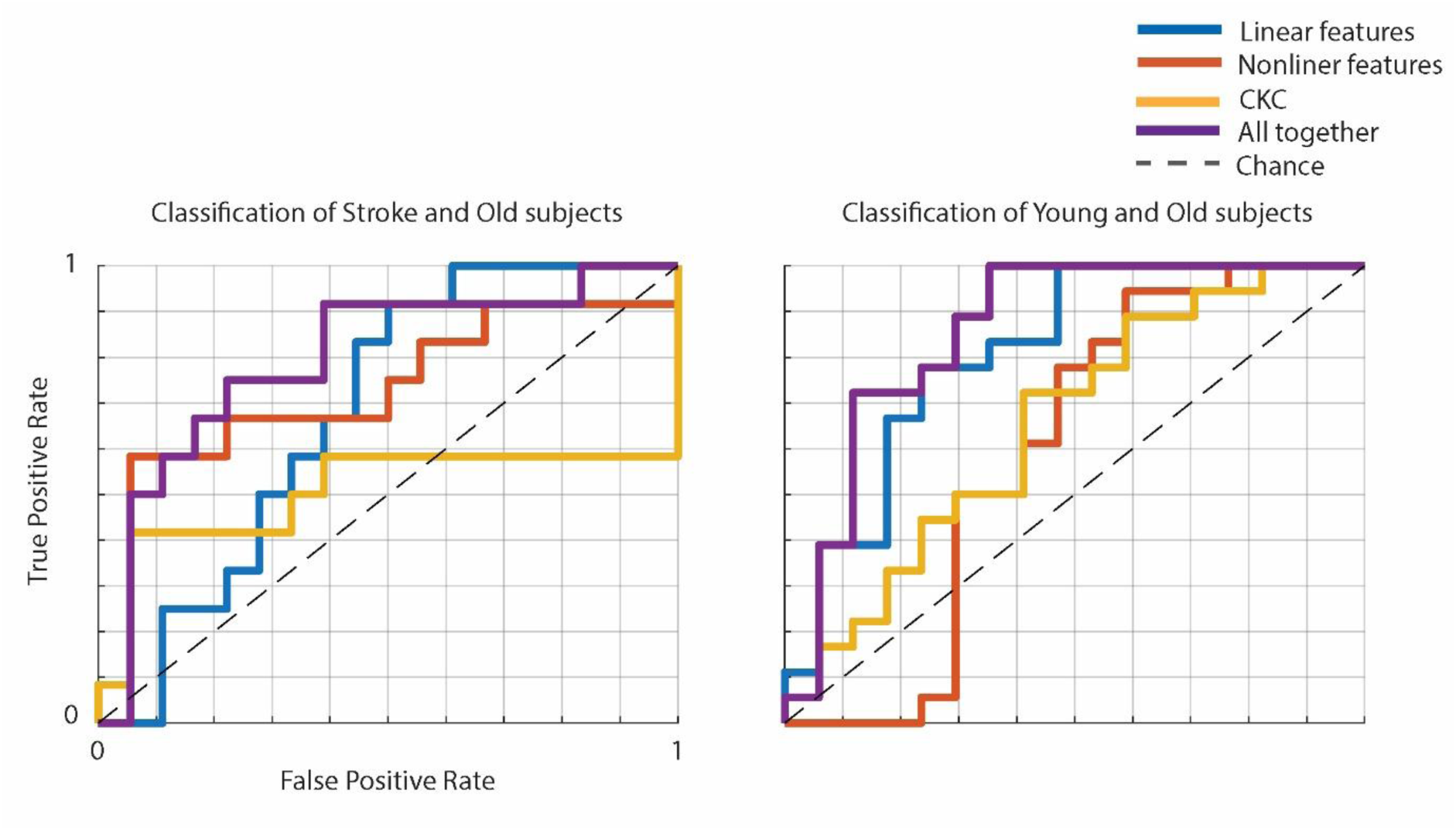
ROC curves for classification of stroke versus older healthy subjects. (**left**) and older versus young subjects (**right**). Models were evaluated using subject-wise leave-one-out cross-validation with logistic regression classifiers. Curves represent performance using linear features (**blue**), nonlinear features (**red**), CKC (**yellow**), and their combination (**purple**). The dashed line indicates chance level.

**Table 1.**
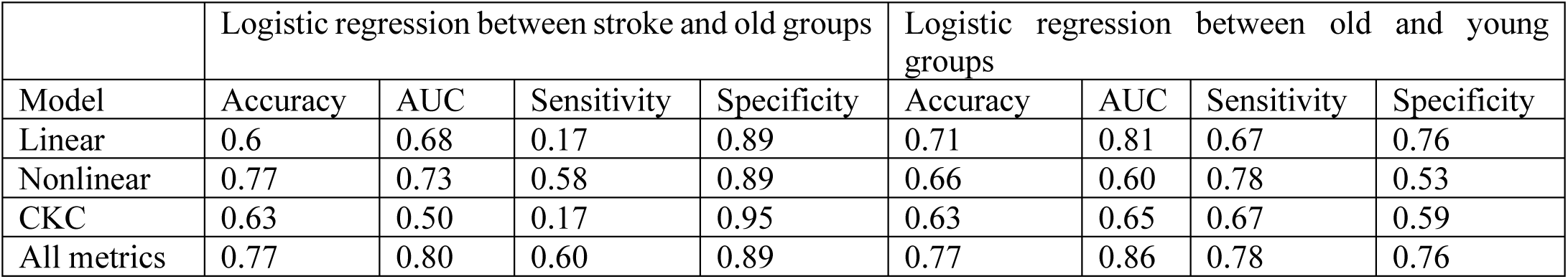
Performance metrics of logistic regression for classification of stroke vs. older healthy subjects as well as older vs. young subjects using subject-wise leave-one-out cross-validation.

|  | Logistic regression between stroke and old groups |  |  |  | Logistic regression between old and young groups |  |  |  |
| --- | --- | --- | --- | --- | --- | --- | --- | --- |
| Model | Accuracy | AUC | Sensitivity | Specificity | Accuracy | AUC | Sensitivity | Specificity |
| Linear | 0.6 | 0.68 | 0.17 | 0.89 | 0.71 | 0.81 | 0.67 | 0.76 |
| Nonlinear | 0.77 | 0.73 | 0.58 | 0.89 | 0.66 | 0.60 | 0.78 | 0.53 |
| CKC | 0.63 | 0.50 | 0.17 | 0.95 | 0.63 | 0.65 | 0.67 | 0.59 |
| All metrics | 0.77 | 0.80 | 0.60 | 0.89 | 0.77 | 0.86 | 0.78 | 0.76 |

Table 1 presents the result of the classification of older healthy versus young subjects. The linear model achieved an accuracy of 71% and an AUC of 0.81, which were the highest accuracy and AUC among the individual feature sets. The combined model further improved performance measures, reaching 77% accuracy and an AUC of 0.86 (Figs 3, Table 1).

In the classification of all three groups, linear features alone provided relatively good performance for distinguishing young subjects (76.5% accuracy) but performed poorly for the stroke group (16% accuracy). Nonlinear features achieved good performance for identifying older subjects (77.8% accuracy) but showed poor performance for the other two groups. CKC features alone resulted in moderate performance for older and young groups (61.1% and 58.8% accuracy, respectively) and again poor performance for the stroke group (25% accuracy). Combining all features substantially improved overall classification metrics, yielding accuracies of 66.7%, 76.5%, and 50.0% for older, young, and stroke subjects, respectively, with the most notable improvement observed in stroke classification (Fig. 4).

**Fig. 4.**
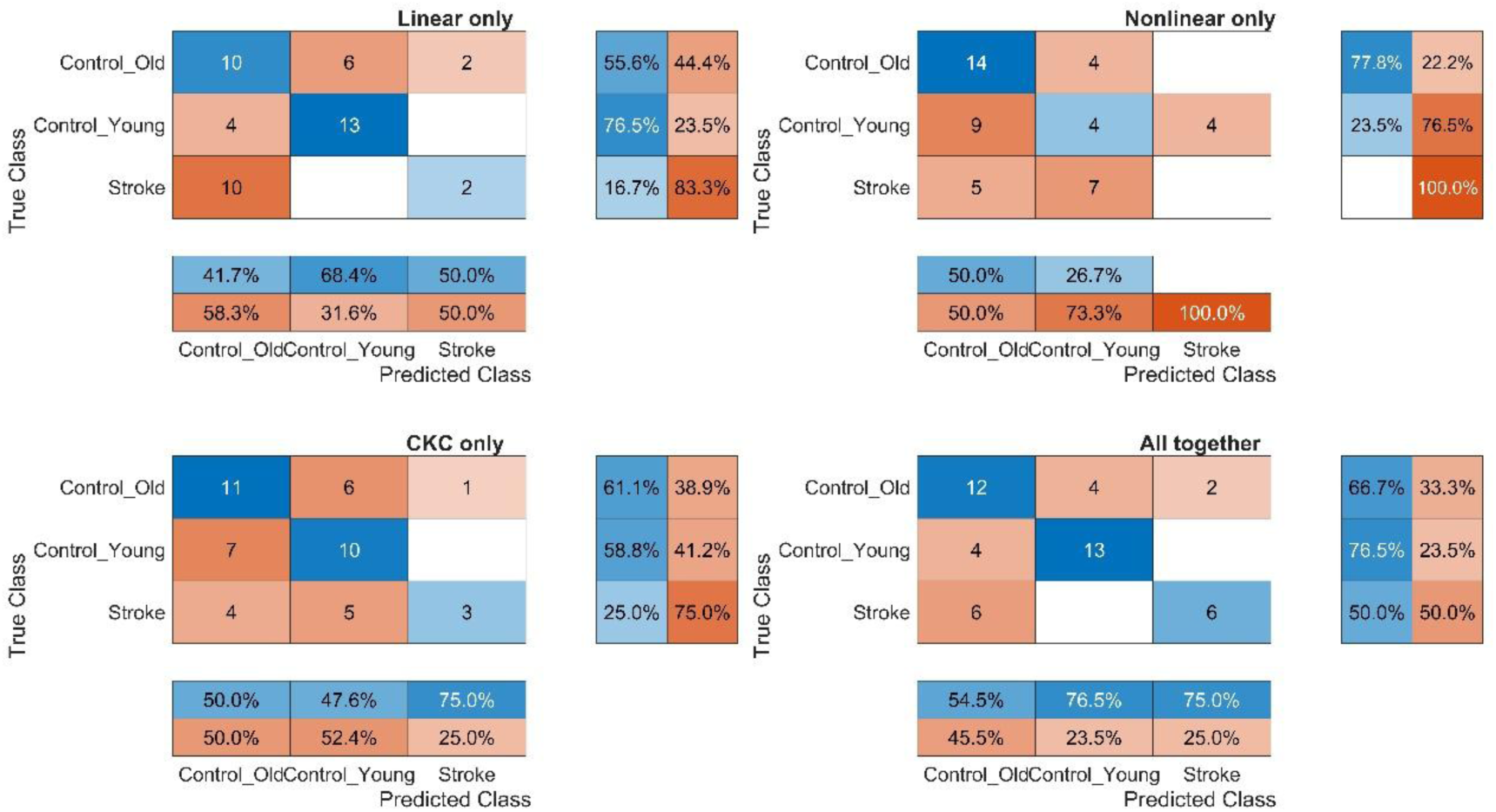
Confusion matrices for subject-wise classification of stroke, old, and young groups with logistic regression. Performance is shown for four feature sets: linear, nonlinear, CKC, and all features combined. Main panels display raw counts, supplemented by row-normalized recall/sensitivity (**right**) and column-normalized precision/positive predictive value (**bottom**).

These results highlight the importance of combining feature types, with nonlinear features contributing most strongly to the discrimination between stroke and older healthy subjects.

### 3.5. State-space transition analysis

Since significant associations between CKC and nonlinear CoP features were observed only in the CoP_ML_ direction, subsequent state-space analyses focused on CoP_ML_-derived variables. Moreover, corresponding analyses based on rCoP-derived variables did not reveal significant effects and are therefore not reported further.

Fig. 5 presents the evolution of brain–body control states across sensory conditions. The first two principal components explained 47.7% and 20% of the variance, respectively. Group-average trajectories across the four sensory conditions revealed distinct patterns of state-space organization between young, older, and stroke participants.

**Fig. 5.**
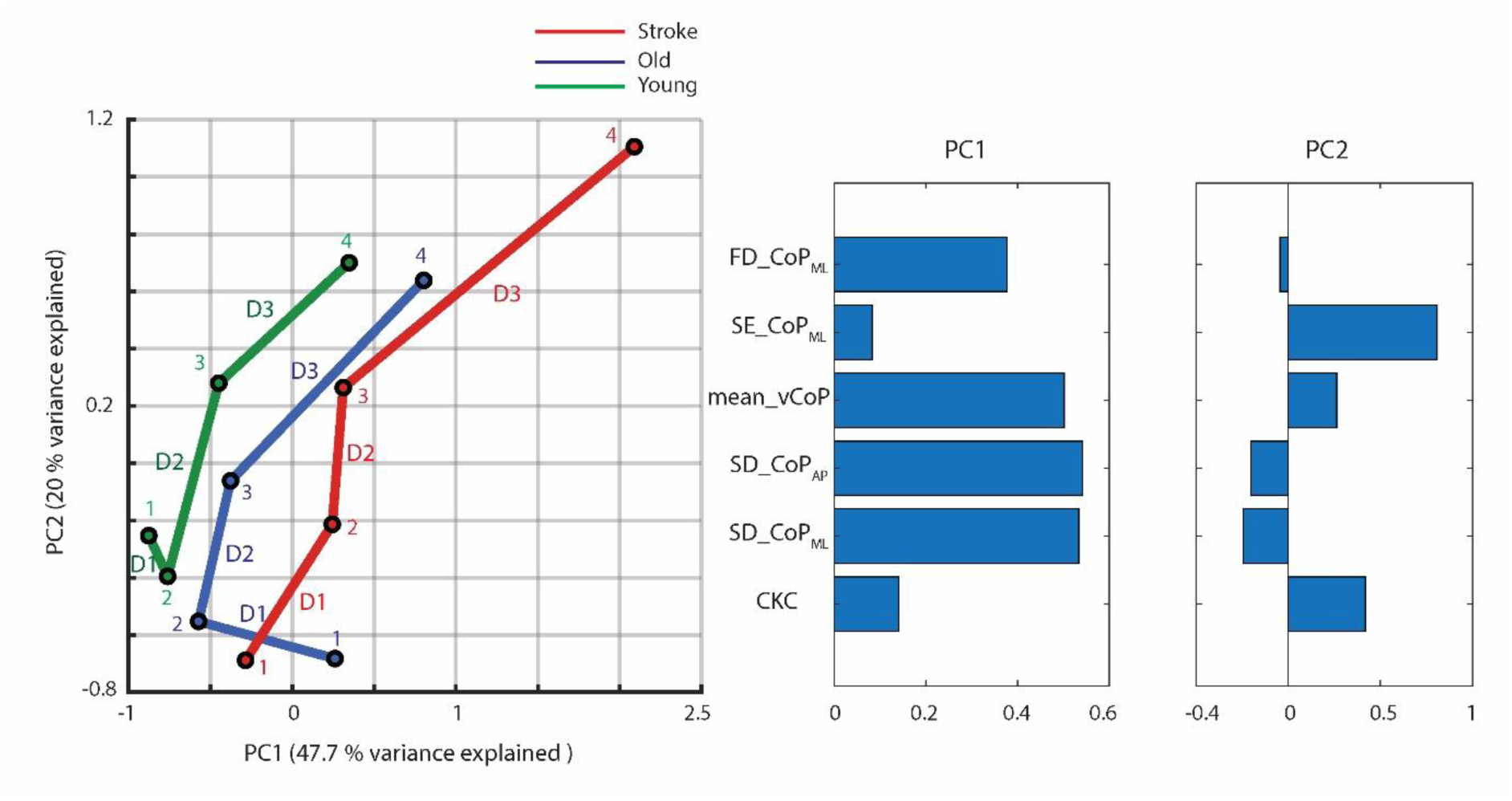
Brain–body state-space organization across sensory conditions for CoP_ML_. Group-average trajectories in the ML brain–body state space projected onto the first two principal components (EOhard = 1, EChard = 2, EOfoam = 3, and ECfoam = 4). Each trajectory represents the evolution of group-average brain–body control states across progressively increasing sensory challenge conditions. The right panels show the PCA weights of the variables for PC1 and PC2.

To quantify transition-specific state-space reconfiguration, Euclidean distances between consecutive sensory conditions were calculated for each participant (D1, D2, and D3).

For the D1 transition (EOhard → EChard), no significant differences were observed between young and older adults (p = 0.65, Cohen’s d = 0.31) or between older adults and stroke participants (p = 0.98, Cohen’s d = –0.19).

For the D2 transition (EChard → EOfoam), no significant differences were observed between young and older adults (p = 0.35, Cohen’s d = –0.35) or between older adults and stroke participants (p = 0.41, Cohen’s d = 0.30).

For the D3 transition (EOfoam → ECfoam), no significant difference was observed between young and older adults (p = 0.09, Cohen’s d = –0.59), while stroke participants exhibited significantly greater D3 transition distances than older adults (p = 0.001, Cohen’s d = 1.19).

Inspection of PCA weights revealed that PC1 was primarily influenced by SD_CoP_ML_, SD_CoP_AP_, mean_vCoP, and FD_CoP_ML_, whereas PC2 was dominated by SE_CoP_ML_ and CKC (Fig. 5).

## 4. Discussion

This study investigated whether nonlinear features of postural sway provide complementary information beyond classical linear measures about sway-based CKC and whether their contribution to CKC differs among stroke survivors, healthy older adults, and young adults. Nonlinear postural sway features demonstrated group– and feature-dependent associations with CKC for CoP_ML_. Importantly, the inclusion of nonlinear features helped explain CKC for CoP_ML_ beyond baseline models containing only linear measures, particularly in the stroke group. Moreover, integrating CKC with both linear and nonlinear CoP features substantially improved group discrimination, reaching up to 77 percent classification accuracy. Notably, the use of multidimensional state vectors constructed from ML-direction linear and nonlinear features and CKC, revealed abnormal reorganization of brain–body control in stroke participants under severe sensory challenge. Specifically, stroke participants showed disproportionately larger state-space transitions during the EOfoam to ECfoam condition change, suggesting altered sensorimotor reconfiguration compared with healthy older adults.

### 4.1. CKC as a Neurophysiological Marker

The present study demonstrated significant associations between CKC and nonlinear features of postural sway in the ML direction in stroke survivors and healthy older adults. These findings suggest that the functional relevance of CKC may extend beyond the strength of brain– body coupling itself and may relate to the temporal structure of postural sway. This interpretation is supported by previous studies demonstrating that CKC is behaviorally relevant across a range of motor and balance-related domains. Specifically, sway-based CKC has been associated with clinical measures of balance and sensorimotor function, including the Berg Balance Scale and Fugl–Meyer Assessment [9]. CKC has also been linked to upright stance stability and to the weighting of vestibular and proprioceptive sensory information during balance control [10]. Moreover, CKC during movement has been shown to relate to motor performance [30]. Previous findings and the present results suggest that CKC may serve as a neurophysiological marker linking cortical activity to behaviorally relevant characteristics of postural control.

### 4.2. Nonlinear dynamics reveal group-specific cortical-postural control strategies

The observed association between CKC and SE in older adults and stroke survivors suggests that cortical activity becomes increasingly linked to the temporal structure of postural sway in these groups. Previous studies have shown that aging and stroke are accompanied by increased cortical involvement during balance control, reflecting a greater reliance on cortical resources to maintain postural stability [9, 10, 31]. In this context, the observed CKC–SE relationship may reflect a compensatory mechanism whereby cortical processes become more closely coupled to the temporal organization of postural sway in order to support balance regulation. Such an interpretation is plausible because SE has been proposed to reflect the adaptability and automaticity of postural control, and balance regulation is known to become less automatic in aging and neurological disorders [32, 33]. Accordingly, the association between CKC and SE may indicate that cortical resources are increasingly linked in regulating the temporal structure of postural fluctuations when maintaining balance requires greater conscious control and supervision.

The observed relationship between FD and CKC suggests that cortical–postural coupling becomes linked to the multiscale geometric organization of ML sway in stroke survivors. Since this association was not observed in either healthy older adults or young adults, it may reflect a characteristic of post-stroke balance control rather than a general feature of aging or standing balance. One possible explanation is that stroke-related impairments, such as asymmetric weight distribution [34], disrupted sensorimotor integration [35], and reduced efficiency of automatic balance-control mechanisms [31], alter the organization of postural sway. Under these conditions, cortical processes may no longer be involved only in generating or correcting postural responses, but may also become continuously engaged in regulating the structure of sway itself. As a result, FD, which characterizes the multiscale geometric organization of postural fluctuations [36], becomes associated with CKC. In this interpretation, the observed FD–CKC relationship may reflect a compensatory mechanism whereby the nervous system establishes a stronger link between cortical activity and the organization of postural sway in order to cope with stroke-related impairments and maintain postural stability.

The predominance of significant associations in the ML direction may reflect the distinct control demands of ML balance regulation. rCoP measures primarily represent the overall magnitude of postural fluctuations by combining multiple sway components into a global measure, while ML sway preserves direction-specific dynamics. Since rCoP incorporates AP sway, which is more strongly influenced by passive biomechanical constraints such as ankle stiffness and inverted-pendulum-like dynamics, it may be less sensitive to cortical contributions underlying active balance regulation [37]. In contrast, ML stability depends more on active sensorimotor regulation and continuous corrective adjustments, making ML sway more sensitive to changes in cortical contribution, particularly when automatic balance control is reduced [38]. In aging and stroke, where sensory integration and postural automaticity are compromised [39], cortical–postural synchronization may therefore become more strongly linked to the temporal and structural organization of ML sway dynamics. This interpretation is also clinically relevant, as ML instability is often more closely associated with impaired balance control and fall risk than AP sway [40].

### 4.3. State-Space Signatures of Postural Control Impairment in Stroke

Beyond the feature-specific associations identified above, the state-space analysis suggests that stroke-related balance impairment involves alterations in the overall organization of brain– body control rather than isolated changes in individual variables. The larger transitions observed in stroke participants under the most demanding sensory condition indicate a less stable control configuration, requiring greater reorganization when sensory information becomes progressively degraded. This may reflect the fact that the brain normally relies on sensing and planning to manage redundancy, uncertainty, noise, delays, nonlinearity, and nonstationarity [41]. In stroke patients, proprioceptive and cross-modal processing deficits [42, 43] and disrupted sensorimotor function [44] impair both sensing and planning, limiting the ability of the balance system to remain within a predictable range. Consequently, the organization of body movements in these individuals under challenging sensory conditions is markedly altered. Although the motor system is thought to plan movements according to a set of principles within a coordinate system whose nature is not fully determined [41], the multidimensional states introduced here provide a practical representation of some of these latent control dimensions. This interpretation is also consistent with the view that motor control operates within a high-dimensional nonlinear state space, where behavior emerges from the interaction of multiple neural and biomechanical processes rather than isolated control variables [45, 46].

### 4.4. Potential biomarkers and clinical implications

As discussed in the previous section, the relationship between CKC and FD was significant only in stroke participants in the ML direction and substantially improved model fit, suggesting that this coupling pattern may reflect stroke-specific alterations in cortical–postural control. In addition, the association between CKC and SE observed in both stroke and older healthy individuals also improved model performance, particularly in the stroke group. These findings suggest that CKC–SE and CKC–FD coupling signatures may represent candidate biomarkers of altered balance control, although further validation in larger and independent populations is required.

Furthermore, the improvement in classification performance obtained by combining linear, nonlinear, and CKC features suggests that each feature type contributes complementary information regarding balance regulation. Specifically, CKC reflects cortical–postural synchronization within the sensorimotor system, linear features capture the magnitude of sway behavior, and nonlinear features characterize the temporal and structural organization of postural control dynamics. The integration of these measures into multidimensional state vectors, which revealed alterations in brain–body state-space trajectories in stroke participants, further supports the importance of combining neural, linear, and nonlinear features to characterize balance impairment.

From a translational perspective, the present findings suggest that CKC–nonlinear feature coupling may provide a framework for identifying deviations from typical brain–body control patterns. For example, in individuals with suspected balance impairment, such as those with recurrent falls or subtle postural complaints, assessing whether CKC–SE or CKC–FD coupling deviates from normative values could help reveal alterations in balance-control strategies that may not be captured by sway magnitude alone. This possibility may be particularly relevant for CKC–SE coupling, which was observed in both older adults and stroke survivors, suggesting sensitivity to altered balance regulation across different levels of impairment. However, this interpretation remains exploratory and requires validation in larger cohorts and in individuals with subclinical or early-stage balance dysfunction.

In addition, the observed group-specific coupling signatures suggest that aging– and stroke-related balance impairments may arise from partially distinct cortical–postural control mechanisms. Consequently, interventions that are effective for one population may not necessarily target the same mechanisms in another, highlighting the need for future studies to investigate whether rehabilitation strategies should be tailored according to specific cortical– postural control profiles. The contribution of this study in this regard is that, given the significant associations emerged predominantly in the ML rather than the rCoP measure, and considering the established relationship between ML instability and fall risk, future rehabilitation approaches may benefit from specifically targeting ML stability and its underlying cortical–postural control mechanisms. Interventions incorporating lateral stability challenges or ML-focused feedback paradigms could provide a useful framework for investigating whether modulation of ML sway dynamics is associated with changes in cortical– postural coupling.

### 4.5. Limitations and future perspectives

Although repeated measurements across multiple sensory conditions and the application of linear mixed-effects modeling have been used in this study, the relatively limited sample size, suggests that the present findings should be considered as an initial proof-of-concept of this framework. Therefore, further studies with larger and independent cohorts are required to confirm the robustness and reproducibility of these findings.

The limited sample size also affects the interpretation of classification performance. Although classification accuracy was evaluated using subject-wise cross-validation to minimize data leakage, external validation using larger independent datasets will be necessary to establish the generalizability and clinical reliability of the proposed framework.

Neural control mechanisms need to be further investigated in future studies. Two possible future directions are the use of perturbation-based paradigms to probe feedback control processes [47], and longitudinal studies examining how structural signatures evolve over time or in response to intervention.

## 5. Conclusion

The present study demonstrates that cortico-postural coupling during standing is associated not only with the magnitude of postural sway but also with its temporal and structural organization.

These associations were group– and feature-dependent, as CKC was associated with SE in both healthy older adults and stroke survivors, whereas its association with FD was observed only in stroke survivors, with no significant associations detected in young adults. These associations emerged predominantly in the ML direction and explained variance in CKC beyond that captured by conventional linear sway measures. Furthermore, multidimensional state-space analysis revealed altered brain–body reorganization across sensory conditions in stroke participants. These findings support the use of nonlinear dynamical features, combined with CKC, as potential markers of balance impairment.

